# Adolescent Peer Victimization Through a Trauma Lens: Heightened Startle Response, Increased Pupil Dilation, Elevated Cortisol, and Reduced Heart Rate Variability in Young Adulthood

**DOI:** 10.1101/2025.05.11.652731

**Authors:** Jens Heumann, Boris B. Quednow, Justin Chumbley, Manuel Eisner, Denis Ribeaud, Lilly Shanahan, Michael J. Shanahan

**Affiliations:** Jacobs Center for Productive Youth Development, University of Zurich, 8050 Zurich, Switzerland; Department of Sociology, University of Zurich, 8050 Zurich, Switzerland; Experimental Pharmacopsychology and Psychological Addiction Research, Department of Psychiatry, Psychotherapy, and Psychosomatics, University Hospital of Psychiatry Zurich, University of Zurich, 8032 Zurich, Switzerland; Neuroscience Center Zurich, ETH Zurich and University of Zurich, 8057 Zurich, Switzerland; Institute of Criminology, University of Cambridge, Cambridge CB3 9DA, United Kingdom; Department of Psychology, University of Zurich, 8050 Zurich, Switzerland

**Keywords:** Peer victimization, Adolescence, Startle response, Autonomic regulation, HPA axis, Stress reactivity

## Abstract

**BACKGROUND:** Adolescent peer victimization (PV) is a prevalent social stressor associated with potential long-term psychological and physiological consequences. Yet, the mechanisms underlying enduring alterations in stress-regulatory systems remain underexplored. This multimodal study examines autonomic, endocrine, and affective reactivity in young adults with a history of severe PV.

**METHODS:** Eyeblink startle electromyograms, pupil dilation, hair cortisol, resting heart rate variability, and self-reported state anxiety were assessed in 182 young adults (age 22) with longitudinal data on adolescent PV (ages 11–20). Effects were estimated using a counterfactual framework with genetically informed inverse probability weighting—leveraging polygenic risk scores and genetic ancestry—and adjustment for time-varying confounders.

**RESULTS:** Victims exhibited heightened startle amplitudes compared to controls, with negative stimuli (angry faces) further exaggerating amplitude and prolonging latency. Pupil dilation increased in response to negative stimuli, but was otherwise blunted. Victims also showed reduced resting heart rate variability, elevated hair cortisol, and slower state anxiety recovery across the session. Depression moderated several effects; more depressed victims showed hypocortisolism and reduced autonomic reactivity.

**CONCLUSIONS:** Findings provide converging evidence and mechanistic insight into potential links between PV and trauma-like physiological effects, suggesting sustained alterations into young adulthood. Targeted interventions are needed to reduce PV and its stress-related dysregulation, preventing potential long-term psychiatric and societal sequelae.

---

Adolescent peer victimization (PV), encompassing verbal and physical violence as well as social exclusion, is a socially pervasive challenge (global prevalence: approximately 30% [1]), with documented mental and somatic outcomes, including anxiety, depression, mood instability, sleeping problems, headaches, and suicidality [2–6]. These outcomes overlap with diagnostic criteria for trauma- and stressor-related disorders, particularly post-traumatic stress disorder (PTSD) and Complex PTSD as defined in the ICD-11 and DSM-5-TR [7–9], suggesting that PV may function as a significant psychosocial stressor that threatens social safety and leads to trauma-like physiological changes [10–12].

While trauma-related physiological impacts are widely studied in PTSD [13–18], their relevance to PV remains poorly understood. Yet one in four children and adolescents develop PTSD after interpersonal trauma [19], with risk arguably influenced by experienced severity [3, 20–22]. Emerging evidence links PV to trauma, with studies identifying associations between victimization and physiological stress markers such as (i) heightened startle response, indicating increased valence-specific defense reactivity [23, 24]; (ii) reduced resting heart-rate variability (RHRV), reflecting diminished vagal control [25, 26]; (iii) blunted cardiovascular responses [27]; and (iv) PTSD symptoms assessed through standardized scales [7, 28–30]. Most studies, however, focus on non-adolescent populations, rely on retrospective self-reports, or examine isolated stress markers, limitations that are addressed by the present study.

Adolescence represents a critical developmental stage marked by heightened stress sensitivity, increased hypothalamic-pituitary-adrenal (HPA) axis reactivity, and maturation of prefrontal regions, including the ‘social brain’ [10, 31–36]. This period reflects a transition from the stress-hyporesponsive phase of childhood to a fully reactive state, with rising cortisol responsiveness that increases vulnerability to HPA axis dysregulation and related psychopathology [10, 37, 38]. Although the long-term effects of childhood adversity on stress physiology are well-documented [3, 5, 39–46], lasting physiological impacts of PV during adolescence remain insufficiently explored.

Indeed, disrupted fear regulation and heightened hyper-vigilance in victimized children [42, 47, 48] point to potential parallels between PV and trauma-like physiological effects. Among key markers, pupil dilation is consistently increased in PTSD and has been proposed as a sensitive index of autonomic hyperarousal following early-life stress [49–53]. Likewise, cumulative cortisol exposure, assessed via hair cortisol concentrations (HCC), reflects long-term HPA axis activity and may index chronic stress load and possible HPA desensitization [10, 54–57]. Notably, these signatures may vary by trauma type and appear further shaped by comorbidities such as depression [58–61].

Effective stress regulation depends on the interplay between parasympathetic activation and HPA axis modulation, processes that are disrupted by trauma [62, 63]. These disruptions impair habituation to repeated stressors [64, 65], potentially compromising emotional and physiological adaptability over time.

Furthermore, stress responses to PV reflect conserved mechanisms, influenced by genetic, psychosocial, and environmental factors [5, 10]. Polygenic risk for PTSD and cortisol-regulating genetic variations have been implicated as key contributors to trauma-induced physiological impacts alongside broader genetic variation [17, 66–68]. Additional factors—including body mass index, physical activity, psychological distress, and substance use—further shape stress responses [69–73].

Considering existing knowledge gaps and previously described complex sources of confounding, this study examines how severe PV changes stress-relevant physiological systems in young adulthood. Using a longitudinal age cohort with a prospectively derived PV classification, we applied genetically informed inverse probability weighting— leveraging polygenic risk scores and genetic ancestry—with additional adjustment for time-varying confounders. The affective startle modulation (ASM) paradigm [74] was used to assess startle eyeblink responses (amplitude and latency) and pupil size throughout the task, including habituation effects. RHRV, HCC, and self-reported state anxiety were also measured to capture autonomic and endocrine regulation. Based on evidence from child maltreatment and PTSD, we anticipated that young adults with a history of severe PV in adolescence (hereafter, victims) would exhibit exaggerated startle response, prolonged latency, increased pupil dilation, reduced RHRV, and elevated HCC—moderated by depression.

## METHODS AND MATERIALS

### Participants

The sample comprised 182 individuals after exclusions (see Procedures) with a mean age of 21.8 years (SD = 0.49); 46.2% were female (*n* = 84), and 11.5% (*n* = 21) were identified as victims. Data were drawn from the Zurich Brain and Immune Gene Study (z-GIG; *n* = 200), a stratified sample of the Zurich Project on the Social Development from Childhood to Adulthood (z-proso) panel (baseline *n ≈* 1,500) [75, 76].This physiological study was part of a larger project examining physiological, biological, fMRI, and psychological correlates of PV in z-GIG.

Severe PV was assessed using longitudinal self-reports from z-proso (ages 11–20). A binary PV classification was derived by contrasting bidirectional peer adversity—covering exclusion, insult, physical violence, and property damage, among other indicators—with severity defined as scoring above the 90th percentile across waves, yielding a conservative classification of victimization based on sustained, uni-directional exposure. Twenty-one individuals (11.1%) were classified as (pure) victims. See Supplement for derivation details.

Startle-eyeblink electromyographic response (EMGR; amplitude and latency), resting heart rate, and state anxiety were assessed in all participants. Sample sizes varied slightly (main: 182, pupil size: 169, RHRV: 154, cortisol: 179), primarily due to technical issues such as equipment malfunctions or incomplete recordings. Demographics were consistent across measures, as detailed in Table S1.

z-GIG study protocols were reviewed and approved by the Zurich Cantonal Ethics Committee (KEK, BASEC Nos. 2017-02083, 2017-02021, and 2023-01951), and written informed consent was obtained from each participant.

### Procedures

During the startle task, participants viewed neutral and angry facial expressions on a computer screen (*≈* 80 cm distance) while exposed to constant 70 dB white noise through Sennheiser HD 202 on-ear headphones. Acoustic startle stimuli were randomly presented while participants viewed expressions. To ensure constant screen distance and minimize head movement, a height-adjustable chin rest was used. The laboratory lacked windows, was continuously illuminated by moderate artificial light, and was maintained at a temperature of 20–23 °C with 55–60% humidity.

The stimuli included 42 faces (21 male, 21 female) from the Chicago Face Database [77], shown with neutral or angry expressions as contrasting treatments. Image luminance was consistent across stimuli (mean = 0.814, SD = 0.030); potential residual differences were accounted for via random slopes for facial expression by subject and a trial intercept. Before each trial, participants fixated on a black dot displayed on a 50% gray screen. Acoustic startle probes (105 dB, 40 ms white noise, < 1 ms rise/fall time) were delivered during two-thirds of the face presentations after randomized intervals of 2 to 5 seconds. Order of faces, startle probe presentations, and delays was randomized once and applied uniformly across participants for consistency.

EMGR was measured using two Kendall H59P electrodes placed on the orbicularis oculi muscle under the right eye, with a reference electrode on the temple. EMGR and stimulus output were recorded via an ADInstruments PowerLab 4/25T system with LabChart 7 software. The setup included an adaptive mains filter, a 50 Hz notch filter, high-pass and low-pass filters at 10 Hz and 200 Hz, respectively, a physical gain setting of 500 µV, and a sampling rate of 1000 Hz. Data post-processing was conducted in R (4.4.3) with custom scripts, ensuring precise signal synchronization, rectification, normalization, blink correction, and baseline correction within the [–50, 10] ms window for consistency and reliability [78, 79]. For analysis, EMGR was quantified in the 20–145 ms window. Trials were excluded for non-identifiable baselines, signal clipping, skin conductance issues (e.g., facial products), or synchronization failures, resulting in 94.9% valid trials.

Pupil size was recorded concurrently during the startle task using an EyeLink 1000 system (v4.51) at a 500 Hz sampling rate, with a monocular desktop mount, head stabilization via the chin rest, and a resolution of 1280 ×1024 pixels. Preprocessing followed standard protocols [80, 81], including the removal of dilation speed outliers, edge artifacts, and deviations from trend lines using R. Blinks were removed and interpolated, followed by smoothing, normalization [(x - min)/(max - min)], and baseline correction within the [–875, 10] ms window. Samples with more than one-third of fixation time outside the facial region were excluded, with the remaining out-of-region time accounted for in the analysis; trials were also excluded due to non-identifiable or shifted baselines, failed blink correction, extreme blinking, resulting in 94.0% valid trials.

ECG was recorded using a Bittium Faros 360 TZ system at a 1 kHz sampling rate. All participants underwent a consistent set of procedures and remained awake throughout the 30-minute measurement period. During ECG recording, they first completed inclusion assessment, followed by a blood draw in most participants and the startle task, with minimal verbal interaction and low physical strain. The first 60 seconds of ECG were removed due to movement-related artifacts. Sneezing and coughing were not specifically tracked, nor were breathing rate or blood pressure; however, given the similar levels of exertion and the overall health of participants, these omissions were considered to have negligible impact on the ECG results.

To assess autonomic function, common RHRV metrics were analyzed [25]: (i) overall RHRV via the standard deviation of all normal-to-normal intervals (SDNN); (ii) long-term variability via the standard deviation of the average normal-to-normal intervals (SDANN) and the standard deviation of successive differences (SDSD); (iii) short-term vagal activity via the root mean square of successive differences (rMSSD) and the proportion of normal-to-normal intervals with successive differences > 50 ms (pNN50); and (iv) sympathovagal balance via the low-to high-frequency ratio ratio (LF/HF).

HCC, available only at age 20, was analyzed from the proximal 3 cm of three thin hair strands collected near the scalp by trained research assistants using liquid chromatography– tandem mass spectrometry (LC–MS/MS), as previously described [82, 83].

State anxiety was measured using the State-Trait Anxiety Inventory (STAI), state subscale [84].

### Polygenic Risk Scores

DNA was isolated from EDTA blood and genotyped using the Illumina Infinium GSA Array, mapped to GRCh38, at LIFE & BRAIN Gmbh, Bonn, Germany, as described in the Supplement. PRSs for PTSD and cortisol-associated degenerative diseases (CDD), adjusting for genetic predispositions influencing stress reactivity, were derived from a multi-ethnic GWAS on trauma susceptibility [85] and a meta-analysis of genetic variants affecting cortisol-binding activity [86]. Quality control included minor allele frequency filtering (MAF > 0.01), imputation quality thresholds (info > 0.8), strand alignment, and sex checks [87]. Summary statistics, originally mapped to GRCh37, were aligned to GRCh38 using the UCSC LiftOver Tool. PRS computations were performed with PLINK 1.90b7 [88, 89] and GNU awk 5.1.0.

### Covariates

Models adjusted for other adversities, including victim-perpetration and sexual harassment/assault, ensuring that the intercept represented individuals without any reported adversity. Principal components (*≈* 75% variance explained) of time-varying confounders (exercise, psychological help-seeking, and substance use) were used to adjust for confounding while minimizing overfitting and preserving statistical power, alongside PRSs for PTSD and CDD, and sine and cosine terms to account for seasonal and diurnal variation. Subjective stress at age 20 and cumulative depression— quantified as the mean across five waves from the SBQ internalizing scale [76, 90, 91]—were also included.

### Statistical Analysis

Data were analyzed in R 4.4.3 using custom scripts. Across all analyses, inverse probability weighting [92], genetically informed by further psychiatric PRSs, genetic ancestry, and biopsychosocial baseline covariates, was applied (gIPW; details in Supplement). The larger control group enabled stable weight estimation and improved comparability with the PV group. Propensity score diagnostics demonstrated successful balancing (maximum absolute standardized difference: 0.198; see Supplement for full diagnostics).

EMGR (amplitude and latency) and pupil dilation were analyzed using mixed models with the established group × image type × trial block (early, late) structure [65, 93, 94]. Random effects accounted for individual and trial-specific variability, with gIPW applied to adjust for confounding. Pupil dilation models included additional adjustments for blink frequency and out-of-region time. State anxiety change was assessed via mixed models comparing pre- and post-experiment measurements using an anxiety × time interaction. Standard errors of mixed-models were bootstrapped with 100K parametric replications to account for pseudo-population variance from gIPW.

RHRV and cortisol (log-transformed) were analyzed using linear models within a complex survey design framework [95], with gIPW weights as sample weights and standard errors from Taylor series linearization [96]. RHRV p-values were FDR-adjusted for multiple comparisons [97]. Cortisol models used corrected cortisol as the outcome, adjusted for hair-related covariates—including color, washing frequency, treatments [56]—by adding residuals from a bootstrap-enhanced lasso regression [98] to the mean, with precision weights based on the inverse variance of corrected cortisol estimates; this approach prevented the increase in estimated parameters. Corrected cortisone was included as a covariate [99, 100].

For each outcome, an additional model was tested including an interaction term PV × depression (PV × DPR).

## RESULTS

Sample descriptives are shown in Table 1, showing no significant baseline differences between the PV and control groups.

**Table 1:**
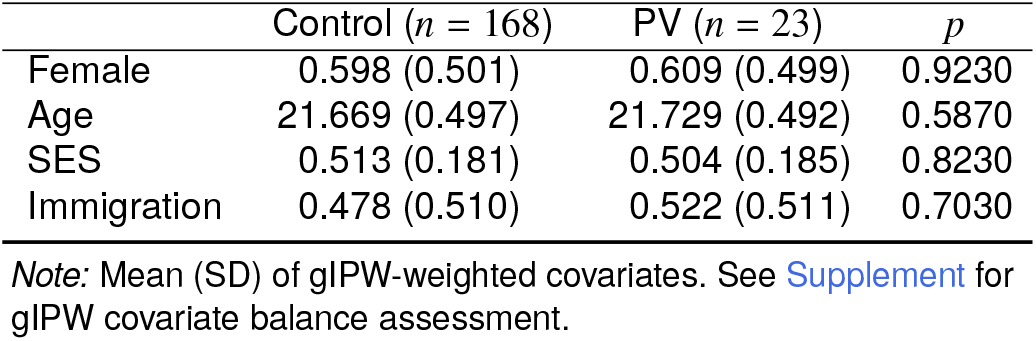
Descriptive Statistics.

| | Control ($n = 168$) | PV ($n = 23$) | $p$ |
| --- | --- | --- | --- |
| Female | 0.598 (0.501) | 0.609 (0.499) | 0.9230 |
| Age | 21.669 (0.497) | 21.729 (0.492) | 0.5870 |
| SES | 0.513 (0.181) | 0.504 (0.185) | 0.8230 |
| Immigration | 0.478 (0.510) | 0.522 (0.511) | 0.7030 |

Note: Mean (SD) of gIPW-weighted covariates. See Supplement for gIPW covariate balance assessment.

### Eyeblink Startle Response

Overall, victims exhibited higher normalized EMGR (mean = 0.135; SEM = 0.008) compared to controls (mean = 0.104; SEM = 0.006). A weighted functional t-test [101] revealed a significant difference in the raw EMGR curves between victims and controls (Mdn *t* = 3.690, Mdn *p* = 0.045), indicating general divergence across the time domain. Over the 20–145 ms window, victims showed a sustained 1.168-fold greater EMGR area under the curve than controls (Fig. 1A).

**Figure 1:**
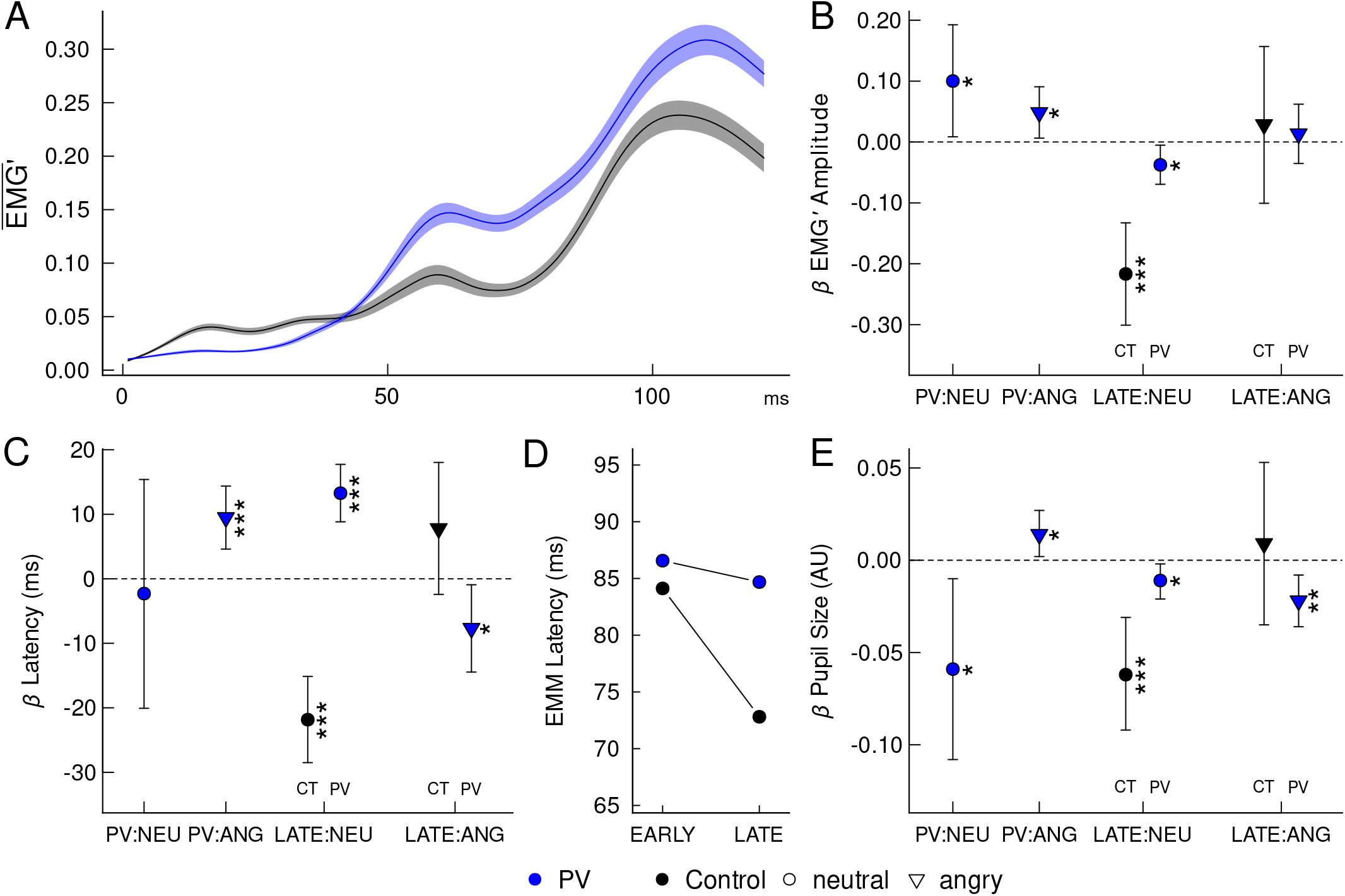
Physiological responses during the startle task. gIPW mixed models with 95% bias-corrected accelerated CIs based on 100K bootstrap reps. **(A)** EMGR curves. Average normalized EMGR response by group, with bands representing SEM. Victims show general divergence across the time domain compared to controls. **(B)** Maximum EMGR. Beta coefficients of maximum normalized EMGR amplitude with 95% CIs. PV = victims, CT = controls, NEU = neutral, ANG = angry, LATE = later trials. Victims’ EMGR was higher with neutral faces and even higher with angry faces compared to controls. During LATE, EMGR decreased for controls viewing neutral faces and was even lower in victims. With angry faces, no additional effects were observed for either group. **(C)** EMGR latencies. Beta coefficients with 95% CIs. ms = milliseconds. With neutral faces, groups showed no significant difference. With angry faces, victims had longer latencies than controls. During LATE, controls’ latencies decreased with neutral faces, while victims’ latencies were longer compared to controls. For angry faces, controls’ latencies tended to be longer in LATE, and victims’ latencies were shorter compared to controls. **(D)** EMMs of latencies (ms). During LATE, latencies decreased for controls, while they essentially stagnated for victims. **(E)** Beta coefficients for pupil dilation in arbitrary units (AU) with 95% CIs. Victims had smaller pupil size with neutral faces compared to controls. With angry faces, victims showed increased pupil size compared to controls. During LATE with neutral faces, pupil size decreased for controls and was even smaller in victims. With angry faces in LATE, pupil size of controls remained unchanged, while victims’ pupil size were smaller compared to controls. \*\*\**p* < 0.001, \*\**p* < 0.01, \**p* < 0.05, ^+^*p* < 0.10

Victims’ maximum normalized EMGR with neutral faces was elevated compared to controls over the 20–145 ms window. With angry faces, victims showed an additional increase in EMGR. In the later trial block (LATE; second half of trials), EMGR decreased in controls when viewing neutral faces and showed a further reduction in victims. For angry faces during LATE, neither controls nor victims showed additional effects (Fig. 1B; Table 2). Adding the interaction term for depression (PV × DPR) did not improve model fit (χ^2^(1) = 0.157, *p* = 0.692).

**Table 2:** Results from Startle Amplitude.

| | $\beta$ | 95% CI | $p$ |
| --- | --- | --- | --- |
| Intercept | 0.431 | [0.291, 0.571] | <0.0001 |
| PV | 0.100 | [0.009, 0.193] | 0.0336 |
| Angry | -0.034 | [-0.128, 0.059] | 0.4771 |
| LATE | -0.217 | [-0.301, -0.133] | <0.0001 |
| Angry × LATE | 0.029 | [-0.101, 0.157] | 0.6631 |
| PV × Angry | 0.048 | [0.006, 0.091] | 0.0246 |
| PV × LATE | -0.038 | [-0.069, -0.005] | 0.0221 |
| PV × Angry × LATE | 0.013 | [-0.035, 0.062] | 0.5922 |

Note: gIPW mixed model. 95% bias-corrected accelerated CIs based on 100K bootstrap reps.

Victims’ EMGR latencies (ms to max. EMGR amplitude) did not significantly differ from controls when viewing neutral faces. With angry faces, however, victims showed longer latencies compared to controls whose latencies did not change significantly. During LATE, latencies decreased for controls when viewing neutral faces but remained longer for victims in comparison. With angry faces, controls’ latencies did not significantly increase during LATE relative to neutral faces, while victims’ latencies were shorter than those of controls (Fig. 1C; Table 3). Estimated marginal means (EMM) revealed that latencies during LATE significantly decreased for controls but essentially stagnated for victims (Fig. 1D). PV × DPR did not improve model fit (χ^2^(1) = 1.019, *p* = 0.313).

**Table 3:** Results from Startle Latency.

| | $\beta$ | 95% CI | $p$ |
| --- | --- | --- | --- |
| Intercept | 70.995 | [45.803, 96.115] | <0.0001 |
| PV | -2.295 | [-20.076, 15.386] | 0.7981 |
| Angry | -3.976 | [-11.409, 3.381] | 0.2907 |
| LATE | -21.839 | [-28.498, -15.130] | <0.0001 |
| Angry × LATE | 7.779 | [-2.421, 18.033] | 0.1360 |
| PV × Angry | 9.484 | [4.607, 14.374] | 0.0001 |
| PV × LATE | 13.268 | [8.829, 17.731] | <0.0001 |
| PV × Angry × LATE | -7.683 | [-14.452, -0.930] | 0.0266 |

Note: gLPW mixed model. 95% bias-corrected accelerated CIs based on 100K bootstrap reps. Angry = angry face, LATE = late trial block.

### Pupil Dilation

Victims’ normalized maximum pupil size over the 20–145 ms startle window was smaller with neutral faces compared to controls. With angry faces, victims exhibited slightly increased pupil size. During LATE, controls’ pupil size decreased when viewing neutral faces and was slightly smaller in victims compared to controls. With angry faces, controls’ pupil size did not show a significant difference during LATE, whereas victims showed a further decrease in pupil size compared to controls (Fig. 1E; Table 4). PV × DPR did not improve model fit (χ^2^(1) = 1.995, *p* = 0.319).

**Table 4:**
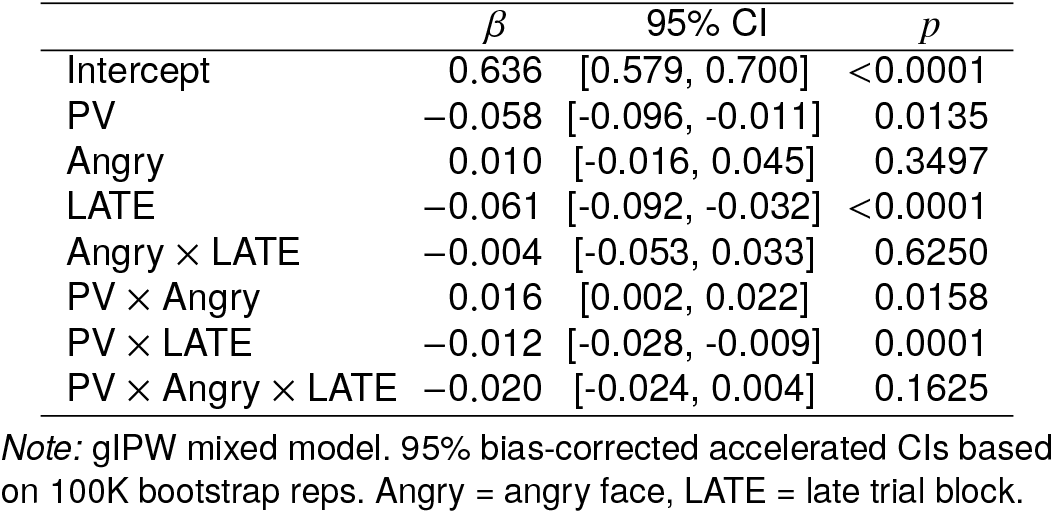
Results from Pupil Size.

### Resting Heart Rate Variability

Victims exhibited consistently lower ln-RHRV across standard time-domain measures—SDNN, SDANN, pNN50, SDSD, and rMSSD—compared to controls. The LF/HF ratio derived from spectral analysis was elevated in victims (Fig. 2A, Table 5). The PV × DPR interaction did not improve model fit for time-domain measures (Mdn χ^2^(1) = 0.269, Mdn *p* = 0.428); however, it did for LF/HF (χ^2^(1) = 0.457, *p* = 0.019), revealing a statistically significant negative interaction with depression.

**Table 5:** Results from RHRV.

| | $\beta$ | 95% CI | $p$ |
| --- | --- | --- | --- |
| PV (SDNN) | -0.308 | [-0.480, -0.136] | 0.0072 |
| PV (SDANN) | -0.285 | [-0.532, -0.039] | 0.0571 |
| PV (pNN50) | -0.494 | [-0.824, -0.163] | 0.0176 |
| PV (SDSD) | -0.508 | [-0.774, -0.241] | 0.0061 |
| PV (rMSSD) | -0.508 | [-0.774, -0.241] | 0.0061 |
| PV (LF/HF) | 0.877 | [0.415, 1.339] | 0.0021 |
| PV × DPR (LF/HF) | -1.168 | [-1.927, -0.410] | 0.0050 |

Note: Table shows coefficients of the PV variable of RHRV models. RHRV outcomes were log-transformed. FDR-adjusted p-values, standard errors from Taylor series linearization. Only in the ln LF/HF model the PV × depression interaction was statistically significant and included in the table. Covariates included but not shown. Variables of interest demonstrated both significance and notable effect sizes.

**Figure 2:**
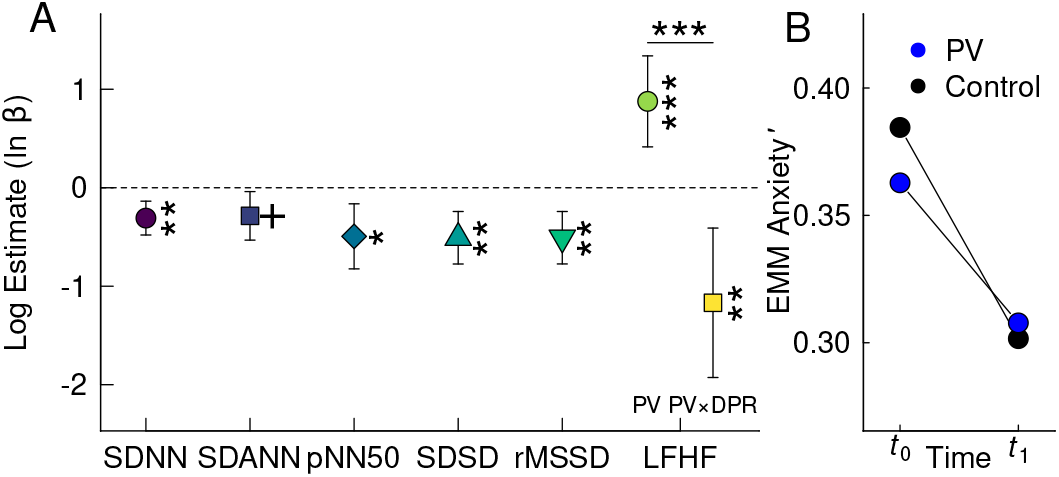
(A) Heart rate variability (RHRV) metrics. ln-transformed *β* coefficients with 95% CIs. Victims showed lower RHRV across SDNN, SDANN, pNN50, SDSD, and rMSSD. LF/HF was higher in non-depressed victims but lower in those with depression. **(B)** Perceived anxiety changes over time. EMMs show perceived anxiety levels at baseline (*t*_0_) and post-treatment (*t*_1_). Victims initially reported slightly lower anxiety compared to controls. Over time, anxiety significantly decreased in controls, while victims showed a smaller reduction, leaving their anxiety levels higher after the experiments. \*\*\**p* < 0.001, \*\**p* < 0.01, \**p* < 0.05, ^+^*p* < 0.10

### Hair Cortisol

On average, cortisol levels were elevated in victims compared to controls. PV × DPR improved model fit (χ^2^ (1) = 23.42, *p* = 0.002) and revealed that cortisol levels were lower in more depressed victims and higher in less depressed ones (Table 6).

**Table 6:**
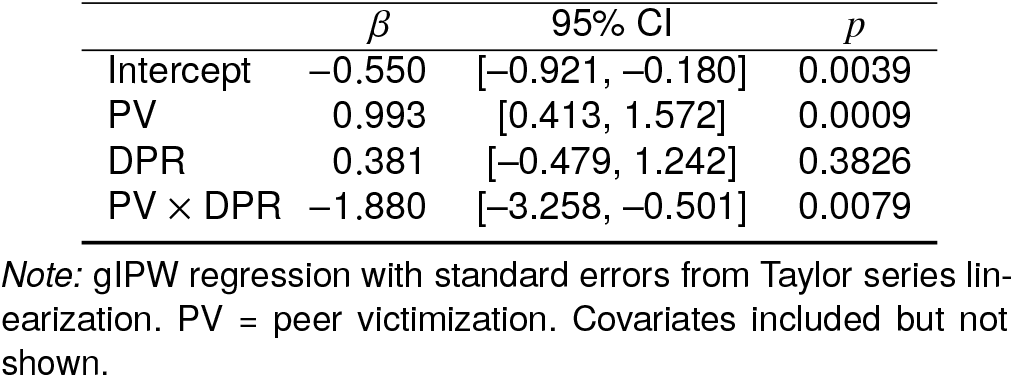
Results from Hair Cortisol.

| | $\beta$ | 95% CI | $p$ |
| --- | --- | --- | --- |
| Intercept | -0.550 | [-0.921, -0.180] | 0.0039 |
| PV | 0.993 | [0.413, 1.572] | 0.0009 |
| DPR | 0.381 | [-0.479, 1.242] | 0.3826 |
| PV × DPR | -1.880 | [-3.258, -0.501] | 0.0079 |

Note: gLPW regression with standard errors from Taylor series linearization. PV = peer victimization. Covariates included but not shown.

### State Anxiety

Victims showed lower state anxiety upon entering the lab compared to controls. Over the course of the experiments, anxiety decreased in both groups. In controls, anxiety decreased considerably, whereas victims exhibited a smaller reduction in anxiety compared to controls, which remained higher after the experimental treatments as indicated by the PV × Time interaction (Fig. 2B, Table 7). Adding the three-way interaction PV × Time × Depression improved model fit (χ^2^(1) = 22.39, *p* < 0.001), but the effect was not statistically significant (*β* = −0.119; 95% CI = [−0.257, 0.019]; *d* = −0.207; *p* = 0.092).

**Table 7:** Results from State Anxiety.

| | $\beta$ | 95% CI | $p$ |
| --- | --- | --- | --- |
| Intercept | 0.373 | [0.333, 0.412] | <0.0001 |
| PV | -0.050 | [-0.087, -0.012] | 0.0095 |
| Time | -0.083 | [-0.095, -0.070] | <0.0001 |
| PV × Time | 0.028 | [0.008, 0.048] | 0.0064 |

Note: gLPW mixed model. Bootstrapped standard errors (100K reps), BCa confidence intervals. Covariates included but not shown. Variables of interest demonstrated both significance and high effect sizes.

## DISCUSSION

This study demonstrates that PV during adolescence is associated with elevated physiological stress susceptibility in young adulthood that mirror trauma-related conditions and, indeed, conditions related to social threat. Using genetically informed inverse probability weighting (gIPW) and a multimodal approach, we identified exaggerated startle, prolonged startle latency, and increased pupil dilation— measured concurrently—in response to negative stimuli. We also observed diminished RHRV, elevated HCC, and slower state anxiety recovery among young adults with a history of severe adolescent PV (victims). These findings converge to suggest that PV constitutes a potent psychosocial stressor with trauma-like physiological effects across systems.

Exaggerated startle responses observed in victims, particularly in reaction to negative stimuli (angry faces) that arguably evoke social threat cues, are consistent with findings from research on childhood adversity [16, 102] and PTSD [103, 104]; exaggerated startle is also a diagnostic criterion for PTSD in both the DSM-5-TR and ICD-11 [8, 9]. This affect-modulation may arise from the brainstem’s role as the central hub of the startle reflex, with neural activation shaped by the affective state of the amygdala [24]. In victims, amygdala activity appears biased toward defensive processing, possibly indicating heightened sensitivity to negative stimuli, as observed in prior cross-sectional research [105]. Decreased startle responses in later trials reflect typical habituation across trials, often attributed to supraspinal modulation and peripheral fatigue [64, 106, 107], with a slightly greater decrease in victims, suggesting increased depletion from their generally exaggerated amplitude.

Victims’ higher startle amplitudes to negative stimuli were accompanied by longer latencies. Amplitude and latency—reflecting response strength and processing speed, respectively—are thought to rely on discordant neural systems [108–110]. In PV, this discordance appears to manifest as exaggerated amplitudes coupled with inhibited latencies, in contrast to facilitated latencies recently observed in PTSD—albeit within a fear-potentiated startle paradigm [111]. While fear learning occurs directly in the amygdala [112], ASM paradigms engage affective context processing, potentially heightened in victims, via slower, higher-order circuits that interact with the amygdala [113]. The subsequent inhibitory influence of victims’ hyperactive amygdala on the brainstem [114] may underlie these prolonged latencies. Such inhibited startle latency has been reported in neuropsychiatric conditions, including psychosis, schizophrenia, Huntington’s disease, and accelerated aging [115–117], suggesting potential long-term consequences for victims. Furthermore, PV during adolescence—a critical period for neural maturation [33, 118]—may disrupt the development of circuits modulating the startle reflex, contributing to latency inhibition [119], which has been linked to larger amygdala volumes [111] consistent with findings of amygdala enlargement in maltreated children [41, 120].

Victims’ reduced pupil sizes may indicate blunted baseline arousal, whereas increased pupil dilation to negative stimuli—paralleling heightened startle amplitude—suggests increased sympathetic arousal, consistent with findings in PTSD [50, 121–123]. Pupil dilation habituation in both groups align with diminishing stimulus novelty [124], with victims’ inhibited dilation in later trials suggesting parasympathetic re-engagement potentially to counteract arousal [49, 51]. Similar attenuation of pupil dilation, observed in social anxiety disorder [125], has been linked to avoidance processes [126], which are also associated with PV [7] and represent a diagnostic criterion for PTSD [8, 9]. Diminished RHRV in victims—reflected in overall (SDNN), long-term (SDANN, SDSD), and short-term (rMSSD, pNN50) measures—and a higher LF/HF ratio align with meta-analytic findings from on PTSD [14, 127]. These patterns indicate reduced vagal activity and total variability at baseline, resulting in a shift toward sympathetic dominance [25, 128]. Conversely, the lower LF/HF in victims with higher depression scores may reflect systemic overdrive beyond homeoregulatory limits, shaped by accumulated somatic and mental load [129–132]. Altogether, such autonomic imbalance aligns with evidence linking chronic social stress to cardiovascular disease [10, 133–135] and emotional dysregulation [136].

Elevated HCC in victims aligns with previous findings from studies on PV and psychosocial stressors more broadly [55, 56, 137–139], reflecting sustained HPA axis activation from chronic social stress. Prolonged activation may blunt immune cell sensitivity to glucocorticoids, resulting in cortisol resistance [57, 140, 141]. Increased cortisol production reflects a persistently altered fight-or-flight response to social threat, often promoting inflammation [142, 143], which has been associated with PV [3, 144]. Moreover, elevated cortisol levels may exacerbate impairments in inhibitory control, as previously discussed, further compounding stress-related effects [138, 145, 146]. In contrast, more depressed victims showed reduced HCC, suggestive of hypocortisolism—a pattern commonly linked to prolonged HPA axis hyperactivity [57, 147–150] and possibly applicable to PV [151]. Hypocortisolism may trigger epigenetic and other molecular processes that contribute to psychopathology, including depression and PTSD [36, 152], with both child maltreatment and PV implicated in long-term depression risk [153, 154].

Finally, victims’ impaired state anxiety attenuation across experimental sessions corroborates physiological evidence of affected inhibitory control, as supported by self-reports. Initially lower state anxiety, combined with a less steep decline compared to controls, aligns with a lack of adaptation to unfamiliar experimental conditions as investigated in PTSD [38]. Alternatively, or interactively, such an adjustment, being age-dependent, may also indicate developmental deficits [155].

The findings reveal a nuanced physiological profile in victims, reflecting recent meta-analytic evidence for the complex etiology of PTSD in youth [156]. Elevated startle amplitudes to neutral stimuli suggest heightened baseline amygdala activity, whereas attenuated pupil dilation reflects blunted arousal. Conversely, exaggerated startle to negative stimuli, accompanied by increased pupil dilation, indicates over-excitation of the amygdala, supported by inhibited latencies and reduced RHRV, suggesting a baseline shift toward sympathetic dominance. This disruption of regulatory balance may reflect overgeneralized avoidance—a hallmark of PTSD—that breaks down under the combined load of startle and negative affective cues, potentially mediated by dysregulated stress-response pathways such as orexin signaling within the amygdalo-cortical-hippocampal circuit [157–160]. During habituation, victims’ reduced startle and pupil responses suggest faster depletion and declining sympathetic dominance, consistent with partial regulatory engagement. The combined heightened arousal and stress reactivity, along with inhibited state anxiety reduction, align with findings linking this profile to elevated PTSD risk [161]. In this context, elevated HCC may reflect sustained dysregulation of the stress system due to repeated activation. Importantly, hypocortisolism and lower LF/HF in depressed victims may indicate that some are already further along a chronic stress dysregulation trajectory, with depression emerging as a downstream manifestation. This pattern may reflect differences in timing (e.g., early vs. late stage), chronicity, and nature (e.g., social exclusion vs. physical violence) of the stressor [151, 162]. These individuals may develop a distinct phenotype characterized by hypocortisolism, with implications for stress-related disorders and chronic conditions such as depression, fatigue, PTSD, pain syndromes, hypertension, fibromyalgia, and other functional somatic disorders [36, 152, 163–165]—also documented in PV and social adversity more broadly [166, 167].

This study has several limitations. First, although PV classification was cautiously derived from comprehensive longitudinal data, the non-PV control group may not represent a fully unaffected baseline. Additionally, victim status was externally classified based on self-reported behaviors and experiences, rather than participants’ self-identification. Second, the sample size limited examination of finer distinctions in PV timing, chronicity, or recency. Third, the propensity score model underlying gIPW was constructed based on genetic data and a thorough literature review, yet residual confounding due to potential model misspecification cannot be excluded. Fourth, although a recognized causal inference method was employed, reverse causation cannot be entirely ruled out, as certain victim characteristics may have contributed to their experiences of PV. Finally, while the Zurich-based sample was population-representative, its high living standard may limit generalizability to broader contexts.

Overall, findings from this genetically informed multimodal study suggest that adolescent PV is associated with alterations in stress physiology in young adulthood resembling trauma-related profiles. These results highlight PV as a clinically relevant form of social adversity with potential long-term biological consequences. Recognizing PV as a modifiable early-life exposure underscores the importance of both prevention and timely intervention to reduce downstream psychological and physiological harm.

## ACKNOWLEDGMENTS AND DISCLOSURES

We gratefully acknowledge the voluntary participation of all z-GIG and z-proso study participants, as well as the support of research assistants involved in data collection.

Funding was provided by the Swiss National Science Foundation (Grants 10531C-197964 to MJS, 405240-69025, 100013 116829, 100014 132124, 100014 149979, 10FI14 - 170409), the Jacobs Foundation (Grants 2010-888, 2013-1081-1), the Jacobs Center for Productive Youth Development, the Swiss Federal Office of Public Health (Grants 2.001391, 8.000665), the Canton of Zurich’s Department of Education, the Swiss Federal Commission on Migration (Grants 03-901 (IMES), E-05-1076), the Julius Baer Foundation, and the Visana Foundation.

Jens Heumann was a fellow of the International Max Planck Research School on the Life Course (LIFE, http:///www.imprs-life.mpg.de) during the preparation of this work.

The authors declare no financial interests or conflicts of interest.

## SUPPLEMENTARY INFORMATION

**Table S1:** Main demographic characteristics of the sample across measures to demonstrate consistency despite slight variations in sample sizes.

|  | Startle | PDL | ANX | HCC | RHRV |
| --- | --- | --- | --- | --- | --- |
| n | 182 | 169 | 182 | 179 | 154 |
| Female (%) | 45.8 | 46.3 | 46.2 | 46.9 | 46.1 |
| Age, mean (SE) | 21.8 (0.49) | 21.8 (0.50) | 21.8 (0.49) | 21.8 (0.50) | 21.8 (0.47) |
| PV (%) | 11.1 | 11.4 | 11.5 | 11.7 | 11.7 |

PDL=pupil dilation, ANX=anxiety, HCC=hair cortisol concentration, RHRV=heart rate variability sample.

### PEER VICTIMIZATION CLASSIFICATION

Rather than relying on direct response counts, PV classification was derived systematically from bidirectional peer adversity data to distinguish victims from non-victims. This approach accounted for the complexities of measuring PV through surveys, where interpreting response patterns was necessary given that self-reports may not directly reflect victimization. Several challenges had to be addressed: (i) Peer adversity was measured at two relational levels—subordinate experiences and dominant behavior that arguably confound each other—using consecutive question blocks across the domains of exclusion, insult, physical violence, and property damage. As z-proso questions only concerned incidence (e.g., “was hit” or “hit someone”), this could indicate victimization, perpetration, or mutual conflict involving differing situational appraisals [1–3]. This ambiguity motivated the definition of victims as those showing consistent subordinate experiences without dominant behavior; (ii) In each wave, peer adversity was assessed retrospectively over the prior 12 months, yielding prospectively derived data with temporal gaps that limited insight into continuous exposure (Fig. S1); (iii) certain forms of victimization and perpetration including sexual adversity, were not consistently assessed across waves, while other unmeasured forms—such as dating or online adversity—may have occurred but were not captured, introducing potential bias from missing information; and (iv) the severity of peer adversity declined over time, necessitating consideration of these downward trends when modeling PV.

**Figure S1:**
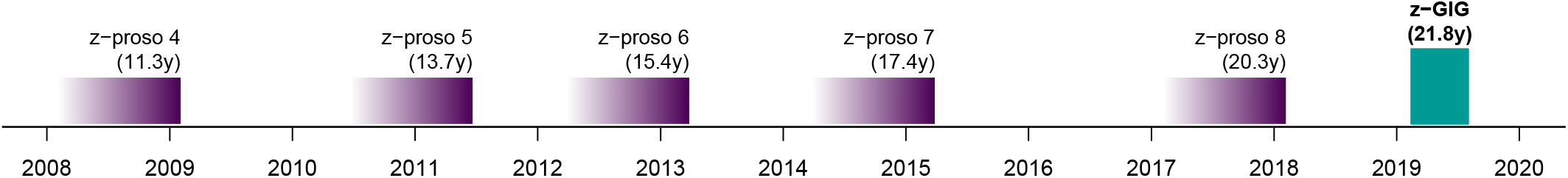
Study timeline illustrating peer adversity assessments in the z-proso panel study. The right edge of each z-proso bar marks a survey wave, with the gradient leftward indicating the 12-month recall period. Gaps denote intervals without data collection.

To address these challenges, individuals who consistently reported high levels of subordinate experiences without reporting dominant behavior were classified as (pure) victims: Incidence was measured using Likert scales (1 = never, 2 = 1 time, 3 = 3 times, 4 = about once a month, 5 = about once a week, 6 = about every day), summed up across domains, and normalized within each relational level to produce composite scores. Participants were considered severely affected on either relational level in a particular wave if they scored at or above the 90th percentile within that wave. Percentile thresholds reflected the observed decline in adversity over time (subordinate experiences: 2009 = 0.36, 2011 = 0.36, 2013 = 0.28, 2015 = 0.20, 2018 = 0.20; dominant behavior: 2009 = 0.24, 2011 = 0.36, 2013 = 0.28, 2015 = 0.20, 2018 = 0.16), unlike fixed Likert cutoffs, which would not have adjusted for this trend. By contrasting subordinate experiences with dominant behavior, participants with a net excess of subordinate experiences were classified as victims and coded as a binary variable (Fig. S2). Among those not classified as victims, all 161 participants constituted the control group represented by the model intercept. This group comprised 85 individuals (52.8%) with no observed peer adversity in the assessed domains (violence, insult, property damage, exclusion), 31 (19.3%) who reported primarily dominant behavior, and 45 (28.0%) who reported both subordinate experiences and dominant behavior. These two behavioral patterns were included as covariates to account for potential confounding within peer adversity. Sexual adversity was modeled separately due to its inconsistent assessment and distinct mechanisms.

This analytic structure was designed to adjust for both known and potentially unmeasured confounding, including adversity not captured by our classification, such as online or dating-related experiences, and to improve upon standard approaches by distinguishing unidirectional victimization from mutual or ambiguous encounters while accounting for developmental trends.

### DNA

153 z-GIG samples with sufficient DNA were processed at LIFE & BRAIN GmbH, Bonn, Germany, for high-throughput bulk analysis. DNA was isolated from EDTA blood following the method previously described [4]. Isolated DNA was genotyped using the Illumina Infinium GSA Array mapped to GRCh38. PED files were subjected to basic QC (maf *>* 0.01; info *>* 0.8), strand flipping, sex check, and clumping [5]. PRS for PTSD and CDD, adjusting for genetic predispositions influencing stress reactivity, were derived from a multi-ethnic GWAS on trauma susceptibility [6] and a meta-analysis of genetic variants affecting cortisol-binding activity [7]. PRS included in gIPW were calculated from summary statistics of recognized GWAS in European populations: ADHD [8]; PD [9]; GAD [10]; and MDD [11]. Summary statistics, originally mapped to GRCh37, were converted to GRCh38 using the UCSC LiftOver Tool to ensure alignment with our data. Principal components of genotyping were calculated, with both analyses conducted using PLINK 1.90b7 [12] and GNU awk 5.1.0. DNA data were available for 84.1% of the participants; missing PRS and ancestry principal components were imputed using random forest (missRanger 2.6.0; [13]) based on gIPW covariates. Imputation of biological data was necessary to address the systematic nature of missingness (NMAR), as PV was significantly associated with refusal to provide a blood sample (*β* = −0.109; 95% CI [−0.120, −0.070]; *p* < 0.0001).

**Figure S2:**
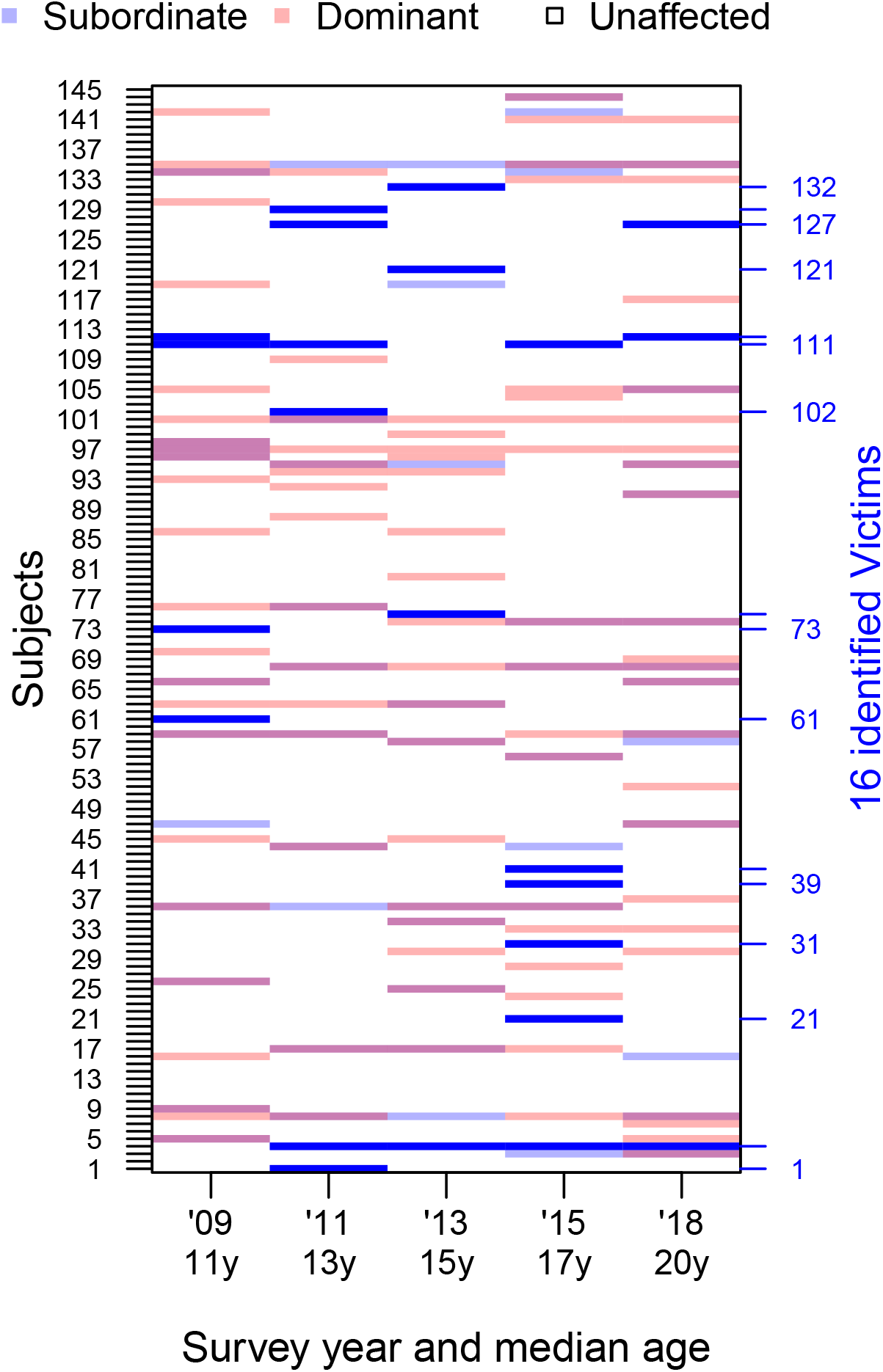
Longitudinal peer adversity profiles. Each row represents a participant, with subordinate (blue) and dominant (purple) experiences shown across survey waves (ages 11–20). Individuals exposed exclusively to recurrent subordinate experiences were classified as peer-victimized (n = 16, marked in bold), forming the victim group at age 22.

### GENETICALLY INFORMED INVERSE PROBABILITY WEIGHTING

Across all analyses, inverse probability weighting [14] was applied to account for both treatment (PV) selection bias and stratified sampling. This counterfactual approach generates a pseudo-population approximating a randomized trial and aids causal interpretation and is generally recommended for observational studies, especially in PV research [15]. To further reduce confounding, weights were genetically informed (gIPW) by psychiatric polygenic risk scores (PRSs), genetic ancestry, and biopsychosocial baseline confounders. Confounders were defined as influencing both PV and outcome [16]. Weights for estimating the average treatment effect on the treated (ATT, Eq. 1) were derived from propensity scores for PV calculated via a logit model,

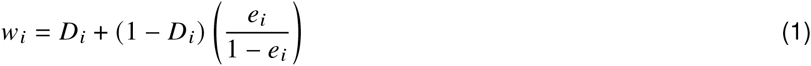

where *D* denotes the PV dummy, and *e* the propensity score [17]. The ATT was chosen as actual victims represent the primary population of interest. To reduce the influence of extreme values, weights were trimmed at the 5th and 95th percentiles. Baseline data for the propensity score model were taken from: (i) self-reports (adolescents, teachers, and parents) from the first three z-proso waves (ages 7–9) prior to the measurement of PV; (ii) PRS for attention-deficit/hyperactivity disorder (ADHD), generalized anxiety disorder (GAD), major depressive disorder (MD), and panic disorder (PD); and (iii) first four principal components from genotyping based on eigenvalue analysis, which showed a clear inflection point at the fourth component with an eigenvalue near 1 (Fig. S3) [18, 19] to account for population structure within the sample (see this Supplement: DNA). Missing values (0.5–6% across all variables) were imputed using random forests (missRanger 2.6.0; [13]). A full list of confounders and corresponding references is provided in Table S2.

**Figure S3:**
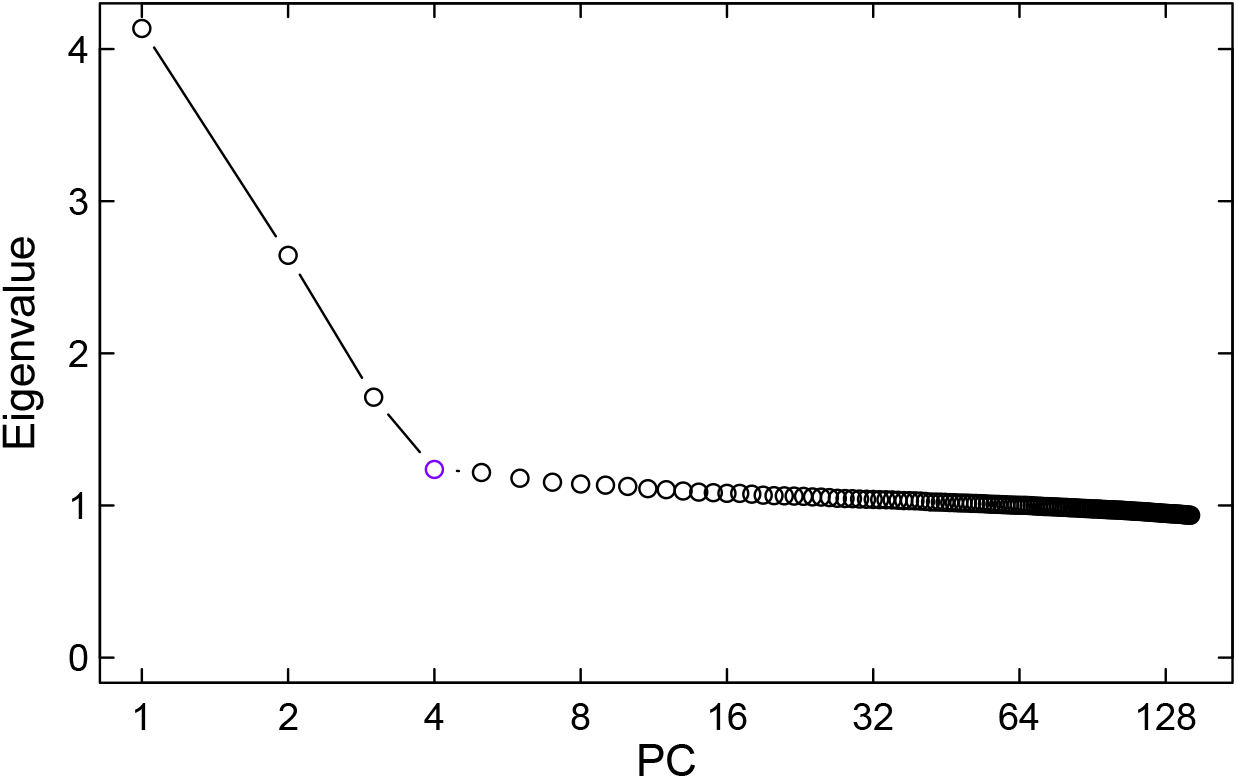
Scree plot of genotyping-derived eigenvalues. The elbow point at component 4 indicates retained ancestry-relevant dimensions.

To minimize extrapolation bias, ATT estimation was restricted to victims with propensity scores within the range of the controls (common support) and excluded three of the original 24 victims due to lack of appropriate matches [20, 21], resulting in the final group of 21 victims. Using complex survey designs [22, 23], gIPW weights *w* were applied as sampling weights throughout the analyses.

**Figure S4:**
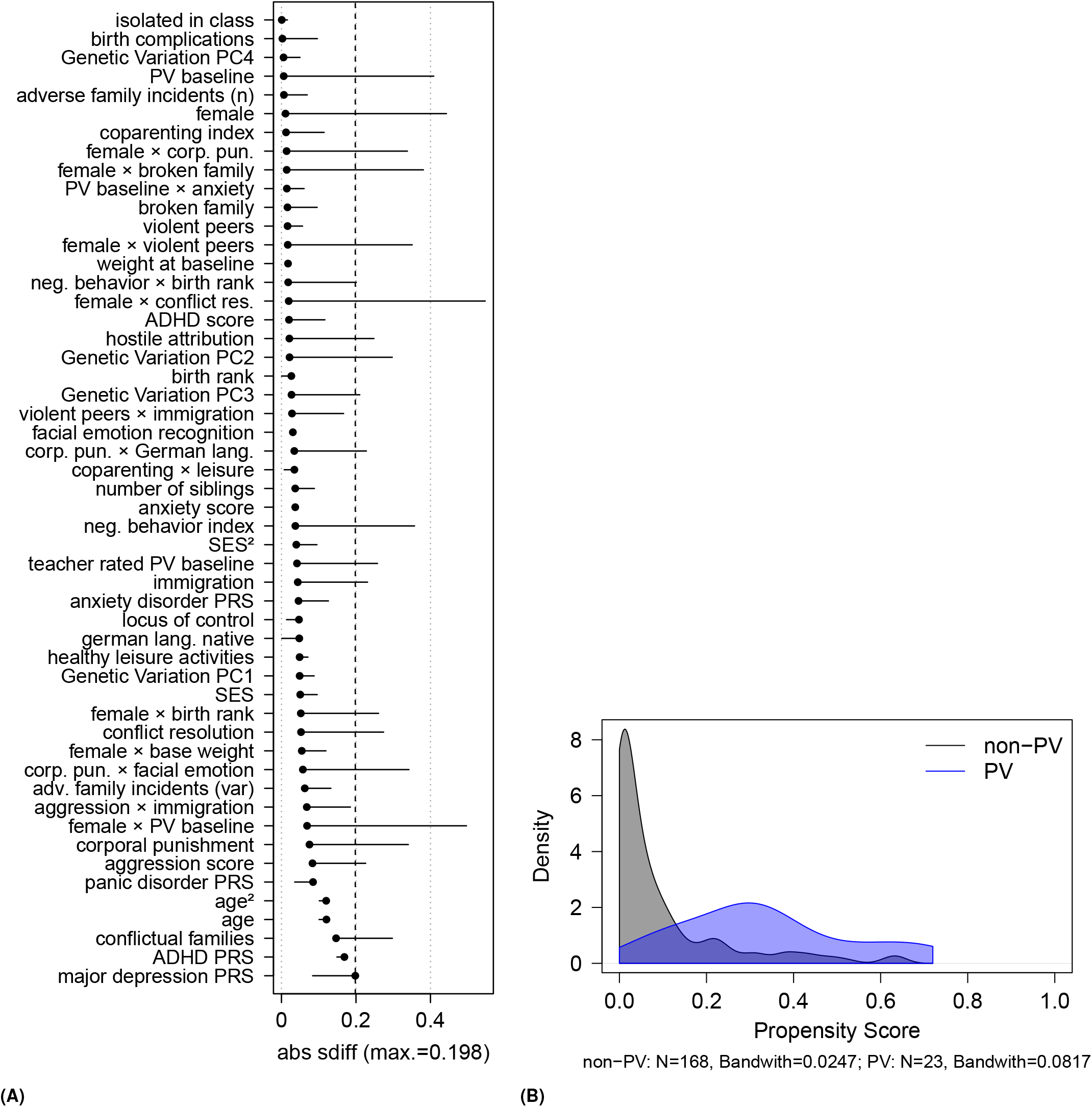
(**A**) Covariate balance before and after gIPW. Absolute standardized differences (ASD) are shown for each covariate used in weighting. Vertical dashed line marks the maximum ASD after weighting (0.198), indicating successful covariate balancing. (**B**) Propensity score distributions. Histogram of propensity scores for treated and control groups in the final analytic sample, indicating sufficient overlap for group comparability.

**Table S2:** Confounders relevant to the relationship between PV and health outcomes, identified through expert knowledge and literature.

| Confounder |
| --- |
| ADHD-score, conflict resolution, corporal punishment, isolated in class, teacher rated bullying [24–26] |
| Adverse family incidents, broken family, conflictual family [27, 28] |
| Age [29] |
| Aggression, anxiety<sup>a</sup>[30, 31] |
| Birth complications [32] |
| Birth rank [33] |
| Coparenting<sup>b</sup>[34] |
| Depression<sup>a</sup>[35, 36] |
| Facial emotion recognition<sup>c</sup>[37] |
| Genetic disposition<sup>d</sup>[38] |
| Healthy leisure activities<sup>e</sup>, body weight (at baseline) [39] |
| Hostile attribution [40] |
| Immigration, German language native [41] |
| Negative behavior in childhood<sup>e</sup>, self-control [24, 42–44] |
| Number of siblings [45, 46] |
| Sex [31, 47, 48] |
| Socio-economic status (SES)<sup>f</sup>[49–52] |
| Violent peers [53] |

Notes: <sup>a</sup>For anxiety and depression, PRS were used, in addition to a score for anxiety. <sup>b</sup>Coparenting score was derived from the extent to which father and mother agreed in their responses to survey questions on parenting. <sup>c</sup>For facial emotion recognition accuracy, a score was created from correct answers to the Assessing Children's Emotional Skills (ACES; [54]) instrument used in z-proso. <sup>d</sup>Genetic dispositions were assessed via PRS derived from GWAS. <sup>e</sup>Healthy leisure and negative behavior scores were created from corresponding z-proso questions. <sup>f</sup>Socio-economic status (SES) was created from parents' income, education, and job prestige (ISEI). References for other scales and instruments can be found in the z-proso project handbook [55].

